# Inferring species interactions from co-occurrence networks with environmental DNA metabarcoding data in a coastal marine food-web

**DOI:** 10.1101/2024.04.24.590890

**Authors:** Elizabeth Boyse, Kevin P. Robinson, Ian M. Carr, Maria Beger, Simon J. Goodman

## Abstract

Improved understanding of biotic interactions is necessary to accurately predict the vulnerability of ecosystems to climate change. Recently, co-occurrence networks built from environmental DNA (eDNA) metabarcoding data have been advocated as a means to explore interspecific interactions in ecological communities exposed to different human and environmental pressures. Co-occurrence networks have been widely used to characterise microbial communities, but it is unclear if they are effective for characterising eukaryotic ecosystems, or whether biotic interactions drive inferred co-occurrences. Here, we assess spatiotemporal variability in the structure and complexity of a North Sea coastal ecosystem inferred from co-occurrence networks and food webs using 60 eDNA samples covering vertebrates and other eukaryotes. We compare topological characteristics and identify potential keystone species, *i.e*., highly connected species, across spatial and temporal subsets, to evaluate variance in community composition and structure. We find consistent trends in topological characteristics across co-occurrence networks and food webs, despite trophic interactions forming a minority of significant co-occurrences. Known keystone species in food webs were not highly connected in co-occurrence networks. The lack of significant trophic interactions detected in co-occurrence networks may result from ecological complexities such as generalist predators having flexible interactions or behavioural partitioning, as well as methodological limitations such as the inability to distinguish age class with eDNA, or co-occurrences being driven by other interaction types or shared environmental requirements. Deriving biotic interactions with co-occurrence networks constructed from eDNA requires further validation in well-understood ecosystems, and improved reporting of methodological limitations, such as species detection uncertainties, which could influence inferred ecosystem complexity.

## Introduction

Quantifying biotic interactions is key to predicting the vulnerability of species and their ecosystems to perturbations from climate change (Foden et al., 2019; Paquette & Hargreaves, 2021). Species distributions are typically assumed to be bound by physiological limits to abiotic conditions at broad scales, while biotic interactions play a secondary role at finer spatial scales (Araújo & Rozenfeld, 2014; Cazelles et al., 2016). However, increasing evidence suggests that certain interactions, *i.e.*, mutualistic or commensalism, can impact species distributions at broad scales and may be especially important for determining species warm-edge range limits (Araújo & Rozenfeld, 2014; Paquette & Hargreaves, 2021). Climate change is disrupting species interactions due to contrasting rates of individual species range shifts and altered phenologies, affecting the timing of interactions and potentially accelerating the loss of species and linked ecosystem functions (Foden et al., 2019; Valiente-Banuet et al., 2015). Assessing the vulnerability of a species to climate change, or predicting their future distributions without accounting for biotic interactions is therefore likely to yield inaccurate outcomes, particularly for specialist species that strongly depend on another species (Foden et al., 2019; Tylianakis et al., 2010). Species interactions are numerous, making it challenging to document all of them directly at whole ecosystem scales, so species co-occurrences have been used as proxies for potential interactions (Morales-Castilla et al., 2015).

The construction of co-occurrence networks is data intensive, requiring community composition data across spatiotemporal gradients (Russo et al., 2022). Environmental DNA (eDNA) metabarcoding data, which provides snapshots of whole community composition within a single sample, can therefore enhance the use of co-occurrence networks (Barroso-Bergada et al., 2021). Interspecific co-occurrences may be indicative of a biotic interaction whereby two interacting species will affect the presence and/or abundance of each other resulting in non-random co-occurrence across space and time (Freilich et al., 2018). Topological characteristics of networks can influence the relative stability or resilience of an ecosystem to perturbations (Barroso-Bergada et al., 2021). For example, high modularity, *i.e.*, separate compartments of non-overlapping, strongly interacting species within the network, may enhance community stability, as the effects of perturbation will be contained within a single module (Delmas et al., 2019). Co-occurrence networks can also identify possible keystone species which have disproportionate negative effects on the whole ecosystem when removed. These species are generally highly connected, with high closeness centrality, *i.e.*, short paths between the node and all other nodes in the network, and betweenness centrality, *i.e.*, node forms shortest paths to connect other nodes (Zamkovaya et al., 2021). However, the proportion of biotic interactions, and which type of interactions contribute to co-occurrence networks is often uncertain, and the interactions detected can vary greatly among replicate networks, even from the same environment (Barroso-Bergada et al., 2021; Russo et al., 2022).

Detecting biotic interactions in co-occurrence networks depends on the type and strength of the interaction, as well as the spatial scale of sampling (Blanchet et al., 2020; Morales-Castilla et al., 2015). Correlation coefficients assume symmetric relationships, but lots of biotic interactions are asymmetric. For example, predation is positive for the predator and negative for prey, whilst the association strength for each species involved in a symmetric interaction, such as mutualism (positive for both species) may differ, potentially obscuring any co-occurrence signal (Blanchet et al., 2020; Goberna & Verdú, 2022). Moreover, the higher the number of interactions per species, the weaker the interaction strength, as the species is less dependent on any one other species (Cazelles et al., 2016). Further, co-occurrences may also stem from shared environmental requirements, dispersal limitations or response to another species, *i.e.*, predator avoidance (Freilich et al., 2018). Indirect interactions can also generate interspecies co-occurrence, whereby a predator may co-occur indirectly with a primary producer due to the predator’s prey being dependent on the primary producer (Blanchet et al., 2020). It is thus impossible to distinguish between co-occurrences derived from biotic interactions and other factors without comparing co-occurrences to previously established biotic interactions. However, to date, co-occurrence networks have largely been assembled for microbial communities, where limited knowledge of functional roles limits validation of the nature of co-occurrences (Berry & Widder, 2014).

Trophic interactions are generally better understood than other interaction types and have subsequently been used most frequently to identify biotic interactions in co-occurrence networks (Ford & Roberts, 2019). Current estimates suggest between a quarter and half of co-occurrences in networks could be trophic in origin. However, these have been primarily estimated using putative trophic interactions for plankton data, which are not well resolved, or using presence-absence datasets (Freilich et al., 2018; Russo et al., 2023; Russo et al., 2022). Detecting trophic interactions in co-occurrence networks is particularly challenging as they exhibit strong spatial dependency resulting in either positive or negative co-occurrences (Cazelles et al., 2016; Russo et al., 2023; Thurman et al., 2019). Negative co-occurrences are expected at finer scales where prey are successfully avoiding predators, whilst positive co-occurrences over greater scales indicate predators tracking their prey (Thurman et al., 2019). Consequently, further exploration comparing known trophic interactions with co-occurrences is needed to validate whether trophic interactions are likely to be detected, and to determine the spatial influence on these relationships. If co-occurrence networks can successfully detect trophic interactions, this could enhance our knowledge of the spatiotemporal variability of trophic interactions, which is often poorly described relative to overall trophic relationships (Young et al., 2015).

Species interactions in the North Sea ecosystem are well characterised as a result of research to quantify negative impacts from fishing pressure and climate change, and the need to understand the consequences of these top-down and bottom-up pressures (Heath, 2005; Lynam et al., 2017). The North Sea represents a “wasp-waist” system, where a few key forage fish species, *i.e.*, sandeels (*Ammodytes* spp.), herring and sprat (clupeids), exert control over the abundance of predators, *i.e*., marine mammals, predatory fishes, seabirds, through bottom-up interactions, and control the abundance of zooplankton prey through top-down interactions (Fauchald et al., 2011; Lynam et al., 2017; Robinson et al., 2023). The abundance and quality of sandeels, the dominant bait fish in the North Sea, has, however, been declining in recent decades, contributing to failures in seabird breeding (MacDonald et al., 2019; Wanless et al., 2005). Forage-fish predators will respond differently depending on diet and foraging specialisations, as well spatial restrictions, *i.e.*, central place foragers are more sensitive to prey depletions (Engelhard et al., 2014). Forage fish species are also vulnerable to climate change due to their reliance on particular substrates (e.g. for burrows or spawning grounds), limiting their ability to redistribute northwards, with temperature driving key lifecycle stageş *i.e.,* spawning (Frederiksen et al., 2011; Petitgas et al., 2013). These changes could lead to further declines and reduced temporal synchrony between predators and prey, where prey are no longer available to predators when required, such as during seasonal foraging periods or when raising young (Edwards & Richardson, 2004). Accordingly, monitoring temporal changes in forage fish availability remains a priority for understanding potential cascading effects across the North Sea ecosystem.

Within the southern Moray Firth, in the northwestern North Sea, the Southern Trench was recently designated as a marine protected area (MPA) for the protection of coastal minke whales which use these waters as a targeted feeding area during the summer months (NatureScot, 2020; Robinson et al., 2023). Here, we investigate spatiotemporal changes in the complexity and structuring of this ecosystem, employing co-occurrence networks and food webs. We explore changes in complexity spatially, between inshore and offshore environments, and temporally during the minke whale foraging season. We quantify differences in co-occurrences detected between spatial and temporal networks and assess the proportion of significant co-occurrences indicative of trophic interactions by comparing co-occurrences with known trophic interactions from the available literature. We identify potential keystone species, *i.e.*, species with the most interactions, and evaluate whether key components of food webs are detected within the co-occurrence networks. We expect interactions in co-occurrence networks to be more volatile as they can stem from different biotic interactions and shared environmental requirements, potentially resulting in key food web components remaining undetected.

## Materials and methods

### Sample collection and analysis

We employed an eDNA metabarcoding dataset derived from 60 samples collected on four sampling trips during June to October 2021 to assess temporal community change, from the southern Moray Firth (Boyse et al., 2023). Marine vertebrate DNA was amplified using two primer sets, MarVer1 and MarVer3, targeting 12S and 16S rRNA barcode markers respectively (Valsecchi et al., 2021; Valsecchi et al., 2020), as well as eukaryotic DNA with 18S rRNA, to capture zooplankton and other invertebrate taxa (Amaral-Zettler et al., 2009; Sawaya et al., 2019). Sequencing libraries were prepared and sequenced separately at the University of Leeds Genomics Facility, St James Hospital, using an Illumina MiSeq Sequencer with 150 bp paired-end lane for both vertebrate primer sets, and 250 bp paired-end lane for 18S rRNA. The bioinformatics pipeline is described fully in Valsecchi et al. (2020) and can be found at http://www.dna-leeds.co.uk/eDNA/. Following the removal of low-quality sequences, PCR duplicates and chimaeras, we clustered sequences into molecular operational taxonomic units (OTUs) using a 98% threshold of homology to the GenBank sequence at species level for the two vertebrate primers, and a 95% threshold at class level for the 18S primer set. We converted read counts into an OTU-specific index, allowing comparison of within-OTU abundances across all samples (Boyse et al., 2023). Firstly, we converted read counts into proportions, then divided the maximum proportion for each OTU from the proportion at individual sites resulting in an index between 0 and 1 for each OTU (Kelly et al., 2019). For vertebrate OTUs that were present across both primer sets, we built an ensemble OTU index by taking the average across both indices at each site (Djurhuus et al., 2020).

### Co-occurrence network construction

We subset our dataset into groups to account for spatial and temporal trends in community composition for co-occurrence analyses (Boyse et al., 2023), as more similar communities produce more specific co-occurrence networks (Berry & Widder, 2014). We will refer to early season, June to July (34 sites), and late season subsets, August to October (23 sites), as ‘temporal subsets’, and will retain spatial signals in the dataset. Nearshore (13 sites) and offshore (47 sites) subsets will be called ‘spatial subsets’ and preserve temporal patterns in the dataset. Small sample sizes (<20) can affect the reliability of co-occurrence networks to accurately predict interactions, so extra caution must be applied to networks produced with sample numbers below this threshold (Hirano & Takemoto, 2019). For each subset, we only retained OTUs that appeared in at least 25% of the sites thereby removing rare species and reducing erroneous correlations in our dataset (Berry & Widder, 2014).

We assembled individual co-occurrence networks by calculating pairwise co-occurrences between species OTU indexes, a measure of relative abundance, with five different metrics (Pearson and Spearman correlations, Bray-Curtis and Kullback-Leibler dissimilarities, and mutual information) in Cytoscape’s Conet plugin (Faust & Raes, 2016). Edges, *i.e.*, significant co-occurrences between two species, were represented in the network if they were supported by at least two metrics, reducing the likelihood of false positives, with the highest and lowest scoring 500 edges being retained to capture both positive and negative interactions (Faust & Raes, 2016). *P* values were calculated using the ReBoot method which compares the null distribution of correlations, accounting for compositionality, using 100 iterations of method and edge specific renormalised permutations, and 100 iterations of bootstrapped confidence intervals of observed correlations (Faust et al., 2012). *P* values across different metrics were then merged using Brown’s method and corrected for multiple testing with the Benjamini-Hochberg approach (Benjamini & Hochberg, 1995; Brown, 1975).

### Food web construction

We used a meta-web approach to construct food webs for the Moray Firth ecosystem by determining all potential trophic interactions between consumers and resources detected by our eDNA metabarcoding dataset (D’Alessandro & Mariani, 2021). We downloaded diet items for fishes, marine mammals and seabirds from FishBase (https://www.fishbase.se; accessed 14/12/2022) and SeaLifeBase (https://www.sealifebase.ca; accessed 6/12/2022), through the ‘rfishbase’ R package version 4.0.0 (Boettiger et al., 2012). We complemented these data with information from the primary literature, including invertebrate species, using the Google Scholar search engine and search terms “Latin species name” or “common species name” with “feeding”, “diet” or “stomach contents”. For some well-studied species, such as marine mammals or seabirds, we restricted the search to dietary studies within the North Sea. We constructed an edge list describing all possible consumer-resource interactions, and subset the data as described above for co-occurrence networks, removing rare species that were present in less than 25% of samples.

Topological properties for both co-occurrence networks and food webs were subsequently calculated using Cytoscape NetworkAnalyzer (Assenov et al., 2008). Food webs and networks were visualised with the iGraph R package version 1.2.1 (Csardi & Nepusz, 2006). Trophic levels for vertebrate species were assigned from FishBase and SeaLifeBase records. We assigned primary producers (*i.e.,* algae) and fungi to trophic level 1, and all other invertebrate classes to trophic level 2 for the purpose of this study. Significant co-occurrences that represented trophic interactions were inferred from the literature used to build food webs. We identified potential keystone species as those with the highest degree of co-occurrences or interactions from co-occurrence networks and food webs respectively (Berry & Widder, 2014).

## Results

### Moray Firth community composition

We retrieved 6,894,772 sequences assigned to 88 OTUs across both vertebrate primer sets, and 1,469,355 sequences assigned to 36 OTUs for 18S rRNA (Boyse et al., 2023). Over 90% of vertebrate reads belonged to teleost fishes, although mammals, Chondrichthyes and birds were also detected. OTUs with the most abundant read counts included forage fish such as sandeels, clupeids and mackerel (*Scomber scombrus*). Classes from Animalia (48% total reads) and Chromista (41% total reads) contributed relatively equally to overall eukaryotic reads. Copepods from the Maxillopoda Class comprised most of the Animalia reads, while dinoflagellates (Dinophyceae) and diatoms (Bacillariophyceae) both dominated Chromista read counts (Boyse et al., 2023). There was a clear temporal signal in community composition, differing significantly between early and late season samples for vertebrates, and across all four sampling months for eukaryotes (Boyse et al., 2023). Sandeels were more prevalent in the early season while mackerel were more abundant in the late season. Maxillopoda accounted for most reads in the first and last sampling months, while Dinophyceae were more prevalent in the middle sampling months (July and August). For both vertebrates and broader eukaryotes, the nearshore community (< 1000 m from shore) had higher alpha diversity and significantly different beta diversity from communities composed of samples collected further offshore (Boyse et al., 2024). Numerous fish species and eukaryote classes were found exclusively in the nearshore community that are known to be associated with shallow depths.

### Temporal food webs and co-occurrence network subsets

We found six and nine fewer nodes in the early season (June and July) compared to the late season (August to October), and 108 and 172 fewer edges in the early season for co-occurrence networks and food webs respectively (Table 1). Edges in the co-occurrence networks were dominated by positive interactions, representing 85.6% of edges in the early season, and 77.9% in the late season, resulting from OTU copresences (Supplementary Figures 1 & 2). Only 53 (17%) edges and 104 (25%) edges in the co-occurrence networks represented known trophic interactions from our literature search. Potential keystone OTUs, *i.e.*, highly connected OTUs, did not overlap between temporal subsets for the co-occurrence networks and food webs, apart from sandeels, which were a keystone OTU for the late season co-occurrence network and both food web subsets (Table 2). Alternative metrics for keystone OTUs, closeness centrality and betweenness centrality, highlighted similar species (Supplementary Tables 1 & 2). We found a negative correlation between average OTU abundance and degree for the early season co-occurrence network (Pearson, *r* = -0.41, p<0.05), but no pattern for the late season network. For example, sandeels only showed a high degree of edge interactions in the late season when they were less abundant (Figure 1). OTUs with the most edge interactions in food webs were dominated by species which occupied mid trophic levels (2-3), as both consumers and prey within the ecosystem (Figure 1). This included some of the most abundant OTUs detected, such as copepods, mackerel, sandeels, and clupeids.

**Table 1.** Topological characteristics of temporal co-occurrence networks (undirected) and food webs (directed) for early season (June to July) and late season (August to October) Moray Firth subsets.

|  | Co-occurrence networks |  | Food webs |  |
| --- | --- | --- | --- | --- |
|  | Early season | Late season | Early season | Late season |
| Nodes<sup>i</sup> | 54 | 60 | 53 | 62 |
| Edges<sup>ii</sup> | 305 | 413 | 290 | 462 |
| Avg. neighbours<sup>iii</sup> | 11.3 | 13.767 | 10.49 | 14.52 |
| Diameter<sup>iv</sup> | 4 | 4 | 5 | 6 |
| Radius<sup>v</sup> | 3 | 2 | 1 | 1 |
| Characteristic path length<sup>vi</sup> | 2.03 | 1.94 | 1.87 | 1.88 |
| Clustering coefficient<sup>vii</sup> | 0.43 | 0.44 | 0.34 | 0.34 |
| Network density<sup>viii</sup> | 0.21 | 0.23 | 0.11 | 0.12 |

<sup>i</sup>Nodes represent OTUS.
<sup>ii</sup>Edges represent interspecific co-occurrences in co-occurrence networks, or trophic interactions in food webs.
<sup>iii</sup>Average number of neighbours refers to the average number of edges per node.
<sup>iv</sup>The diameter is the maximum length of the shortest path between two nodes.
<sup>v</sup>The radius is the minimum length of the shortest path between two nodes.
<sup>vi</sup>The characteristic path length is the average shortest path length between any two nodes in the network.
<sup>vii</sup>The clustering coefficient represents the average clustering coefficient across all nodes and is a value between 0-1. It represents the proportion of co-occurrences among the neighbours of a node.
<sup>viii</sup>Network density describes the proportion of realised co-occurrences from all potential co-occurrences.

**Table 2.** Potential keystone species, *i.e.*, the ten molecular operational taxonomic units (OTUs) with the highest degree (number of edges), identified in co-occurrence networks and food webs from the early season (June to July) and late season (August to October) Moray Firth subsets. Gradients of blue indicate trophic levels, from light blue (trophic level 1) to dark blue (trophic level 4).

| Co-occurrence networks |  | Food webs |  |
| --- | --- | --- | --- |
| Early season | Late season | Early season | Late season |
| Phaeophyceae | *Oncorhynchus* | Polychaeta | Polychaeta |
| *Spinachia spinachia* | Dinophyceae | Gadidae | Gadidae |
| *Dicentrarchus* | *Taurulus bubalis* | Maxillopoda | Maxillopoda |
| *Symphodus melops* | Ammodytidae | Clupeidae | Bivalvia |
| *Centrolabrus exoletus* | *Gasterosteus aculeatus* | Gastropoda | *Scomber scombrus* |
| *Gasterosteus aculeatus* | *Zoarces* | Ammodytidae | Clupeidae |
| *Taurulus bubalis* | *Larus argentatus* | *Scomber scombrus* | Gastropoda |
| *Pholis gunnellus* | *Anguilla anguilla* | *Pomatoschistus minutus* | Ammodytidae |
| *Chirolophis ascanii* | *Pholis gunnellus* | Pleuronectidae | Pleuronectidae |
| *Salmo* | *Phocoena phocoena* | *Trisopterus esmarkii* | *Pomatoschistus minutus* |

**Figure 1.**
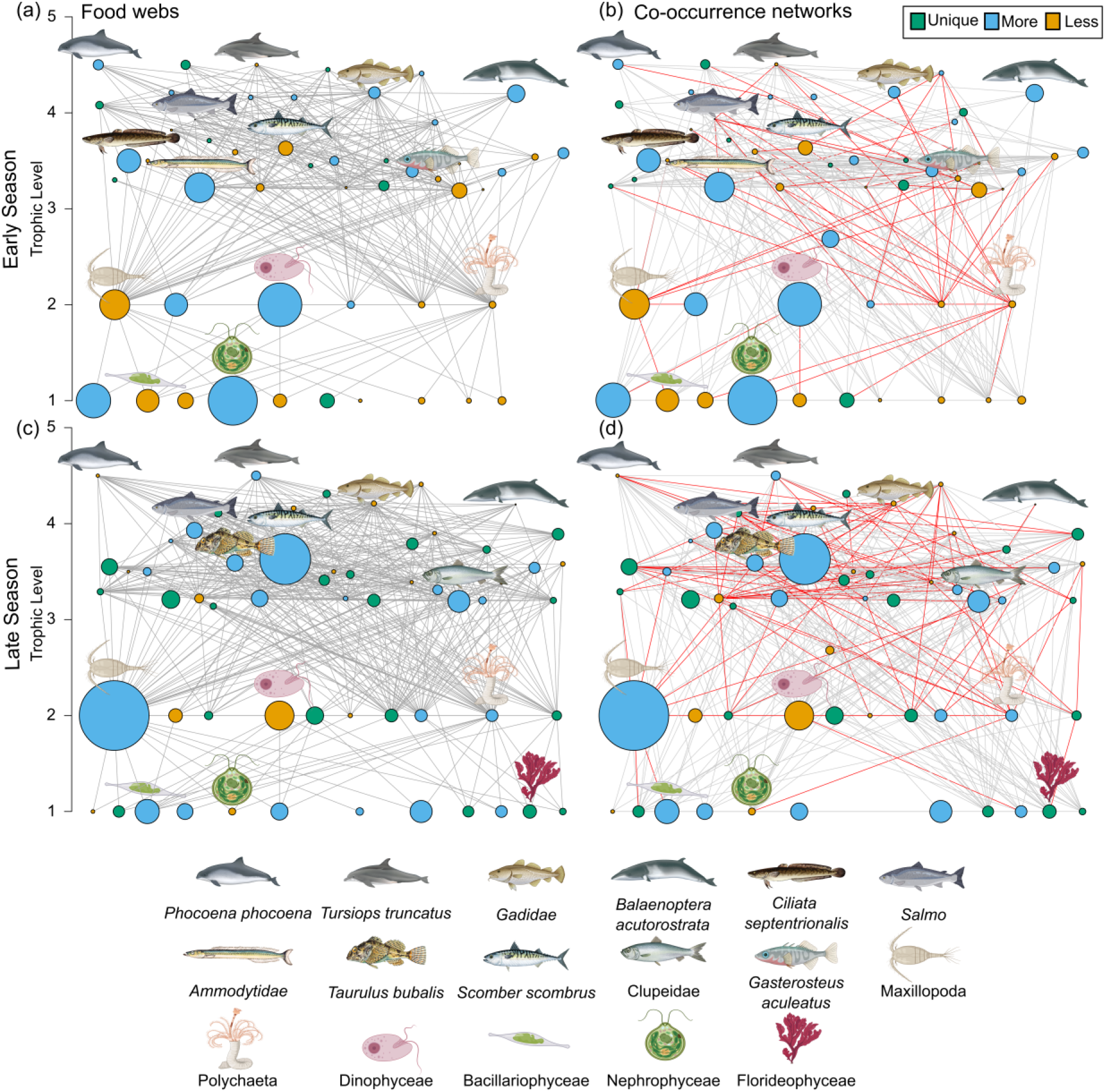
Food webs from (a) early season (June to July) and (c) late season (August to October), determined by environmental DNA metabarcoding detections and known trophic interactions. Respective co-occurrence networks, built with pairwise co-occurrences using 5 different metrics (correlations, dissimilarities, mutual information) and eDNA relative abundance data, for the (b) early season and (d) late season. The size of the node represents the scaled average abundance of the molecular operational taxonomic unit (OTU) across samples, and the colour indicates whether the OTU is unique to that time period (green) or more (blue) or less (yellow) abundant. Red edges in co-occurrence networks signify trophic interactions while grey edges represent all other significant co-occurrences. Individual OTUs are plotted in the same location between graphs.

### Spatial food webs and co-occurrence network subsets

We detected 22 more OTUs in the nearshore community compared to offshore, despite the nearshore community including only 13 samples compared to 47 offshore samples (Table 3). This included 27 OTUs which were only found in the nearshore community (Figure 2). More edges were formed between nodes for the nearshore community, with 139 more edges for the co-occurrence networks, and 206 more edges for the food webs. The spatial subsets also detected a greater proportion of co-presences (nearshore 73.6% and offshore 77.9%), compared to mutual exclusions (Supplementary Figures 3 & 4), and a small proportion of trophic interactions (nearshore 20.9%, offshore 22.5%). Similar to the temporal subsets above, we discovered little overlap between potential keystone OTUs in the co-occurrence networks and food webs, with Gastropoda being the only keystone OTU in both the nearshore co-occurrence networks and food webs, and sandeels the only overlapping keystone OTU offshore (Table 4). Two keystone OTUs, the three-spined stickleback (*Gasterosteus aculeatus*), and the long-spined bullhead (*Taurulus bubalis*), were shared across all four co-occurrence networks. Conversely, the followingt five keystone OTUs were shared by one spatial subset and one temporal subset, dinoflagellates (Dinophyceae), brown algae (Phaephyceae), harbour porpoise (*Phocoena phocoena*) and salmon/trout genuses (*Salmo*, *Oncorhynchus*). We found a positive correlation between average OTU abundance and degree for the nearshore network (Pearson, *r* = 0.31, p<0.05), but no correlation was detected for the offshore network.

**Table 3.** Topological characteristics of spatial co-occurrence networks (undirected) and food webs (directed) for nearshore (<1000 km) and offshore communities.

|  | Co-occurrence networks |  | Food webs |  |
| --- | --- | --- | --- | --- |
|  | Nearshore | Offshore | Nearshore | Offshore |
| Nodes<sup>i</sup> | 68 | 46 | 67 | 46 |
| Edges<sup>ii</sup> | 383 | 244 | 465 | 259 |
| Avg. neighbours<sup>iii</sup> | 11.27 | 10.61 | 13.55 | 10.826 |
| Diameter<sup>iv</sup> | 4 | 3 | 5 | 4 |
| Radius<sup>v</sup> | 3 | 2 | 1 | 1 |
| Characteristic path length<sup>vi</sup> | 2.13 | 1.88 | 1.86 | 1.8 |
| Clustering coefficient<sup>vii</sup> | 0.38 | 0.4 | 0.31 | 0.36 |
| Network density<sup>viii</sup> | 0.17 | 0.24 | 0.11 | 0.13 |

<sup>i</sup>Nodes represent OTUS.
<sup>ii</sup>Edges represent interspecific co-occurrences in co-occurrence networks, or trophic interactions in food webs.
<sup>iii</sup>Average number of neighbours refers to the average number of edges per node.
<sup>iv</sup>The diameter is the maximum length of the shortest path between two nodes.
<sup>v</sup>The radius is the minimum length of the shortest path between two nodes.
<sup>vi</sup>The characteristic path length is the average shortest path length between any two nodes in the network.
<sup>vii</sup>The clustering coefficient represents the average clustering coefficient across all nodes and is a value between 0-1. It represents the proportion of co-occurrences among the neighbours of a node.
<sup>viii</sup>Network density describes the proportion of realised co-occurrences from all potential co-occurrences.

**Table 4.** Potential keystone OTUs, *i.e.*, the ten molecular operational taxonomic units (OTUs) with the highest degree (number of edges), from co-occurrence networks and food webs from the nearshore and offshore communities. Gradients of blue indicate trophic levels, from light blue (trophic level 1) to dark blue (trophic level 4).

| Co-occurrence networks |  | Food webs |  |
| --- | --- | --- | --- |
| Nearshore | Offshore | Nearshore | Offshore |
| *Zoarces* | Dinophyceae | Polychaeta | Polychaeta |
| Calcarea | *Taurulus bubalis* | Maxillopoda | Gadidae |
| Ascidacea | *Limanda limanda* | Gadidae | Maxillopoda |
| Granuloreticulosea | Ammodytidae | Bivalvia | Clupeidae |
| *Taurulus bubalis* | *Salmo* | Gastropoda | Ammodytidae |
| *Phocoena phocoena* | *Gasterosteus aculeatus* | *Scomber scombrus* | *Scomber scombrus* |
| Phaeophyceae | *Oncorhynchus* | Clupeidae | Gastropoda |
| *Gasterosteus aculeatus* | *Tursiops truncatus* | Pleuronectidae | *Pomatoschistus minutus* |
| Gastropoda | Bacillariophyceae | Ammodytidae | Pleuronectidae |
| Chlorodendrophyceae | *Ctenolabrus rupestris* | Ophiuroidea | *Trisopterus esmarkii* |

**Figure 2.**
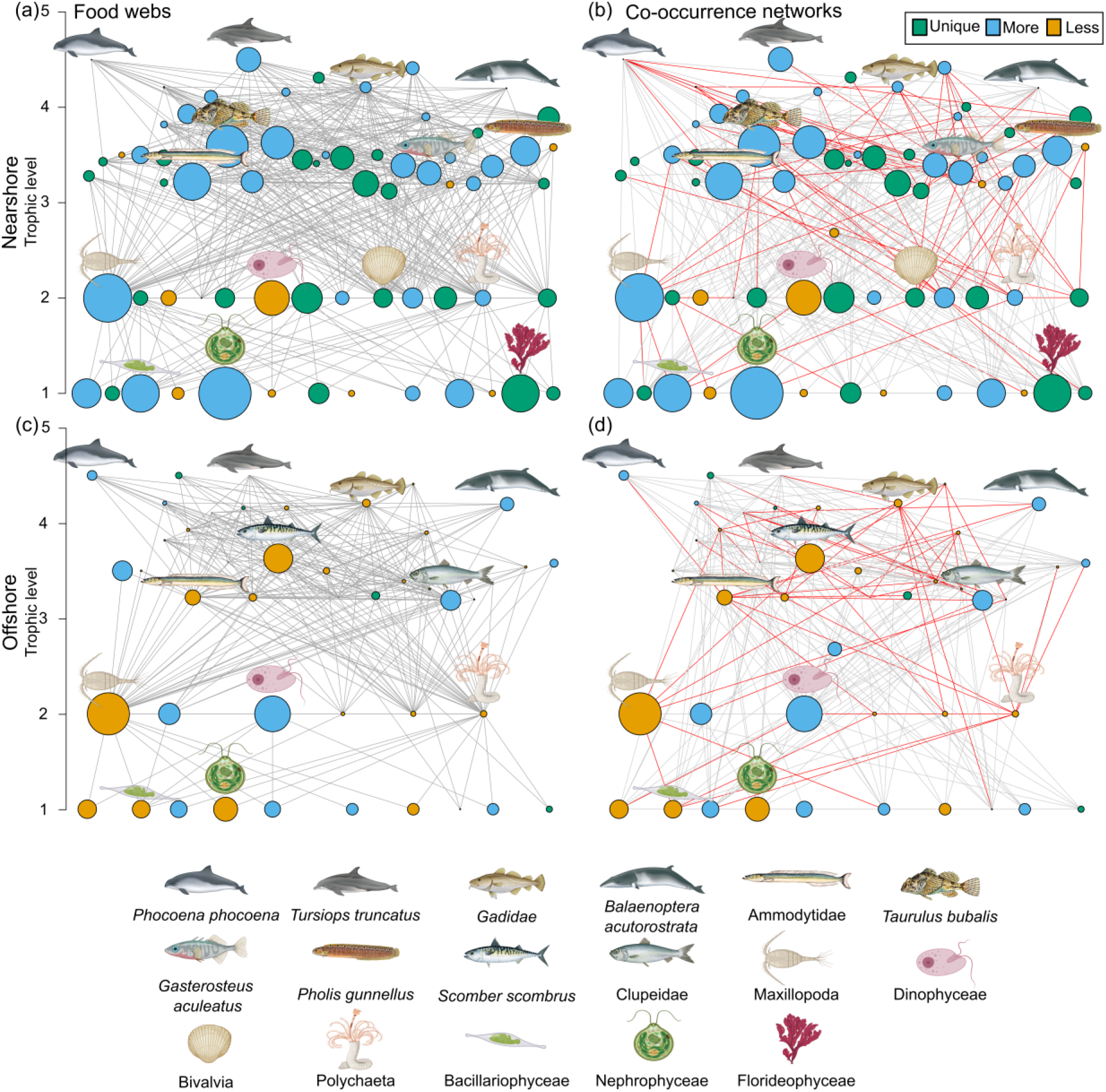
Food webs from the (a) nearshore community with samples collected <1000 m from shore, and (c) offshore community determined by environmental DNA metabarcoding detections and known trophic interactions. Respective co-occurrence networks, built with pairwise co-occurrences using 5 different metrics (correlations, dissimilarities, mutual information) and eDNA relative abundance data, for the (b) nearshore and (d) offshore communities. The size of the node represents the scaled average abundance of the molecular operational taxonomic unit (OTU) across samples, and the colour indicates whether the OTU is unique to that time period (green) or more (blue) or less (yellow) abundant. Red edges in co-occurrence networks signify trophic interactions while grey edges represent all other significant co-occurrences. Individual OTUs are plotted in the same location between graphs.

### Stability of edges between different co-occurrence network subsets

Within the temporal and spatial co-occurrence network subsets, there were a high number of unique edges, with only 61 (9%) and 37 (6%) shared edges in the temporal and spatial subsets respectively (Figure 3a). Despite the temporal and spatial subsets being assembled with the same data, there was a high number of edges only detected in either the temporal or spatial networks. Only 268 edges (27%) were found in both subsets, with a similar number of edges unique to the temporal or spatial subsets, 394 and 325 edges respectively. We investigated edge stability further with cetacean trophic interactions and found very few overlapping edges between subsets (Figure 3b). Only two trophic interactions between bottlenose dolphins (*Tursiops truncatus*) and European seabass (*Dicentrachus*), and between harbour porpoises and sandeels were detected in more than one subset. Only 21% of the detected trophic interactions were with known dominant prey species, and these interactions were all unique to one subset, apart from between harbour porpoises and sandeels which were detected in both nearshore and offshore networks.

**Figure 3.**
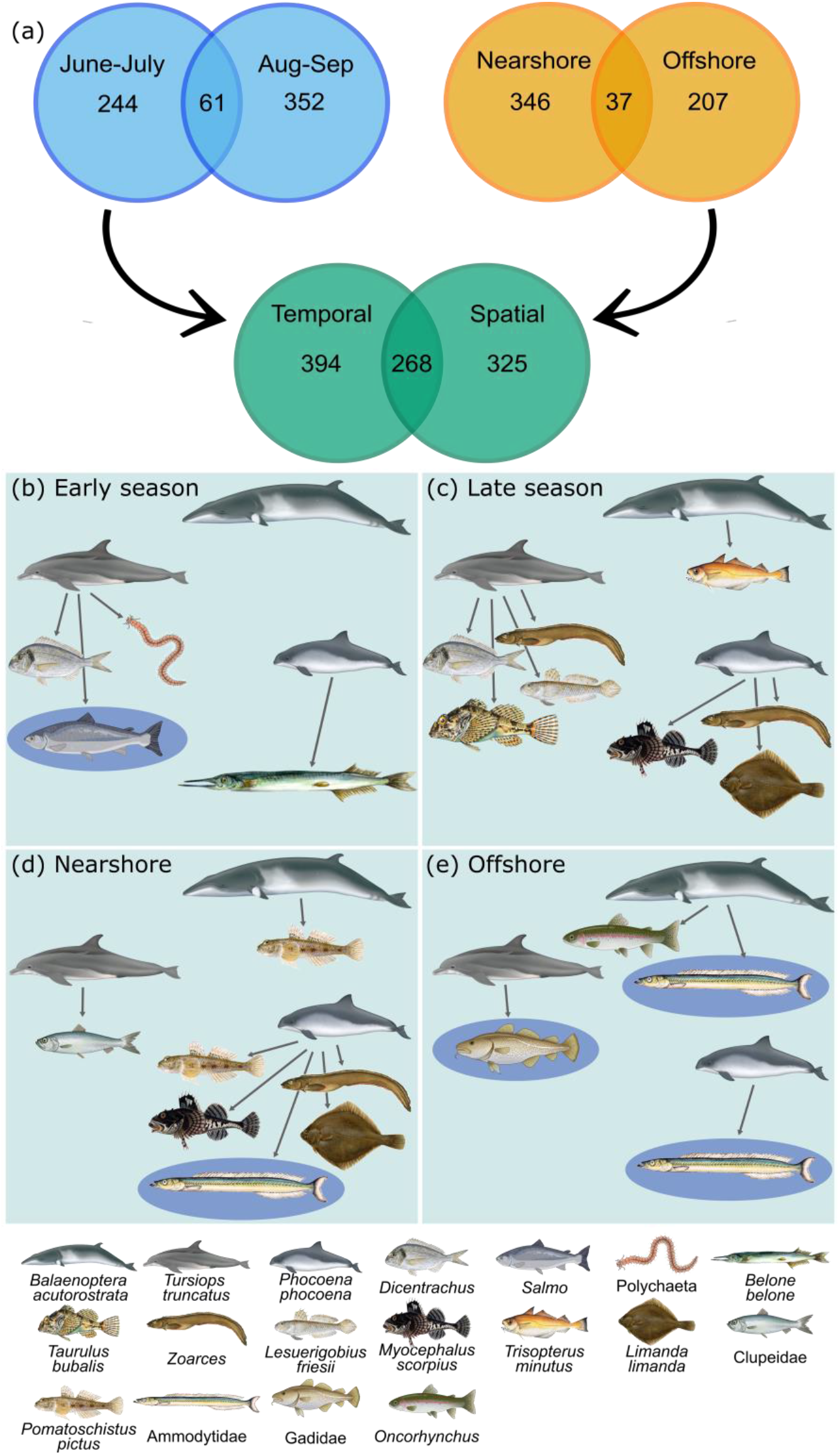
3. (a) Venn diagrams showing the number of overlapping edges detected in the different co-occurrence network subsets. ‘Spatial’ represents all edges in the monthly samples combined and ‘Temporal’ represents all edges in the distance from shore communities combined, with duplicates removed. (b) to (d) show trophic interactions between cetaceans and prey in temporal (early season, late season) and spatial (nearshore, offshore) co-occurrence network subsets. Dark blue ellipses indicate known dominate prey species.

## Discussion

In this study, we assessed spatiotemporal variability in the structure and complexity of the Moray Firth ecosystem and identified potential keystone species derived from food webs and co-occurrence networks. We found consistent trends in topological characteristics used to describe overall co-occurrence networks and food webs, with higher interaction diversity found in the nearshore and late season subsets. However, the interactions contributing to different co-occurrence networks were highly changeable, with only 27% of interactions shared between the spatial and temporal subsets. Significant co-occurrences can stem from different biotic interactions or shared environmental niches, so we hypothesised that trophic interactions would form a minority of co-occurrences. Indeed, we confirmed that trophic interactions only formed a small proportion (<25%) of co-occurrences detected across both our spatial and temporal networks, and significant co-occurrences between cetaceans and their known dominant prey species were rare. As a result, we discovered that keystone OTUs, *i.e.*, highly connected nodes, in food webs were not comparable with those detected in co-occurrence networks.

Topological characteristics did not vary greatly between co-occurrence networks and food webs, or between spatial and temporal subsets (Tables 1 and 3), which is expected given the small spatial scale (tens of kilometres) of sampling in the present study. The nearshore network was produced from fewer samples (13 samples) but shared similar topological characteristics with the other networks, suggesting the small sample size had limited impact on retrieving significant interactions (Hirano & Takemoto, 2019). The most notable differences were found in the number of edges, with higher interaction diversity detected in the late season and nearshore networks. Networks with more interactions could indicate higher ecosystem functionality and greater redundancy of interactions, thus increasing the community’s resilience to disturbance (Tylianakis et al., 2010; Valiente-Banuet et al., 2015). Higher interaction richness in the nearshore subset was likely driven by greater species richness, which has been found to increase closer to shore in previous eDNA metabarcoding studies (Jiménez et al., 2018; O’Donnell et al., 2017; Ríos Castro et al., 2022).

Despite the widespread application of eDNA metabarcoding data for building co-occurrence analyses, methodological limitations impacting topological characteristics have not been addressed sufficiently in these studies to date. For example, the higher species richness of the nearshore environment could be a methodological artefact, since the water samples were collected from surface water (at 4 metres depth; Boyse et al. (2023)), such that more benthic species might be detected in the shallower nearshore samples. Previous studies employing eDNA metabarcoding have detected benthic species at deeper depths (>200 m) due the presence of eddies (O’Donnell et al., 2017), although stratification in the southern Moray firth may prevent DNA mixing in the water column (Adams & Martin, 1986). Notably, our 18S primer set targeting eukaryotes failed to amplify DNA from organisms in the Malacostraca class (Crustacea), such as amphipods, carideans, decapods and isopods. Whilst copepods dominate North Sea plankton assemblages and are the major component of forage fishes’ diets, these other crustacean groups still form part of the diet of most planktivorous forage fish, and therefore would be reasonably well connected in food webs, potentially becoming more important as copepod abundance declines with warming waters (Garzke et al., 2015; Mortelmans et al., 2021; Segers et al., 2007). Further, other potentially important OTUs may have been removed due to incomplete reference libraries, affecting the overall ecosystem complexity, as coverage of North Sea macrofauna with the 18S gene is only approximately 36.4% to date (Sawaya et al., 2019; Zamkovaya et al., 2021). Extracting abundance data from eDNA metabarcoding is debated, especially in relation to methods for tackling bias stemming from differing amplification efficiencies (Hansen et al., 2018; Hestetun et al., 2020; Shelton et al., 2023). Here, we transformed our read counts into an OTU-specific index, which assumes the amplification efficiency is constant across all samples regardless of the community composition (Kelly *et al*., 2019). However, species composition will likely affect the read numbers retrieved, with more accurate abundance estimates recovered for dominant taxa compared with rarer taxa (Skelton et al., 2022). All these effects could potentially impact interaction diversity and network complexity so must therefore be acknowledged as shortfalls when pairing eDNA metabarcoding data with co-occurrence network analyses.

Only a small proportion of co-occurrences were attributable to trophic interactions in the present analyses. Our food webs showed that most predators within the study area feed on multiple prey species (Figures 1 and 2), thus reducing the likelihood of detecting trophic interactions due to dietary plasticity (Robinson et al., 2023; Thurman et al., 2019). Some trophic interactions may not be described in the literature and are subsequently categorised as non-trophic interactions in our analyses. For example, dinoflagellates, which often make up a significant component of plankton biomass and are both important consumers and a food resource, were not well connected in our food webs, likely owing to difficulties identifying planktonic species in stomach content analyses (Pethybridge et al., 2014; Sherr & Sherr, 2007). In this case, co-occurrence network analyses can potentially contribute to our current understanding, as dinoflagellates are an abundant and well-connected OTU in the offshore network, suggesting they may play an important role in this community. Increased diet metabarcoding studies will thus serve to improve our knowledge of trophic interactions in plankton communities in the future (Zamora-Terol et al., 2020).

Detecting trophic relationships in co-occurrence networks relies on the assumption that predators are tracking their prey (co-presence) or prey are avoiding their predators (mutual exclusion) (Thurman et al., 2019). In reality, however, species will partition their time between different behaviours or, at the extreme end of the scale, show seasonality in their foraging and breeding grounds, which will affect interspecific interactions over different spatiotemporal scales (Risch et al., 2014). An important limitation to eDNA sampling, which often overlooked in co-occurrence networks, is that the method is unable to distinguish between age classes, e.g. larval versus adult stages (Hansen et al., 2018). Age substantially influences what a species eats and who it is eaten by (Bossier et al., 2020). For example, herring will consume juvenile cod, but adult cod will also consume herring, which may skew correlation trends when age-class cannot be distinguished (Lynam et al., 2017). Spawning events may additionally result in increased peaks in eDNA abundances which could further bias interactions towards those incorporating early life stages, since the presence of DNA derived from gametes may not reflect the occurrence of trophic interactions in the same way as for older age classes (Di Muri et al., 2022; Valsecchi et al., 2021).

We discovered high numbers of edges unique to either the spatial or temporal subsets, despite the same data being used to create these independent subsets (Figure 3). Designating subsets prior to co-occurrence analyses is often carried-out arbitrarily, but samples must come from similar environments to avoid habitat filtering dominating potential interactions (Berry & Widder, 2014). In this study, we focused on trophic interactions with cetacean species to investigate edge stability, as we know that cetacean diets are typically dominated by a few target prey species, thus increasing the likelihood of detecting the trophic interactions present (Thurman et al., 2019). For example, sandeels and clupeids comprise over 80% by weight of minke whale (*Balaenoptera acutorostrata*) diets in Scottish waters, whilst sandeels and whiting (Gadidae) make up 80% of harbour porpoise diets (Pierce et al., 2004; Santos et al., 2004). However, significant co-occurrences between sandeels and minke whales or harbour porpoises were only detected in the spatial co-occurrence subsets (Figure 3). This may be due to the spatial subsets retaining a temporal signal, and the abundance of sandeels displays strong temporal variability with higher abundances in June and July when they are actively feeding within the water column (Boyse et al., 2023; Henriksen et al., 2021). Similarly, we also detected higher abundances of both minke whales and harbour porpoises in June and July. Gadidae species contribute up to 84% of the bottlenose dolphin diet, and salmon are also suspected to be a dominate prey species, although their otoliths are almost completely digested so are inherently difficult to detect in stomach content analyses (Santos et al., 2001). We found co-occurrences between bottlenose dolphins and Gadidae in the offshore network, and with salmon in the early season network, when species occurred at very low abundances. Zurell et al. (2018) previously observed that predator-prey co-occurrences were more likely to be detected when both species were rare, and detectability decreased as one of the species became more abundant. However, rare species in eDNA analyses may represent false positive detections stemming from transport of DNA in tides and currents (Hansen et al., 2018). All three of these cetacean species also displayed trophic interactions with species not known to form large portions of their diet, which could indicate prey switching, although this seems unlikely as their targeted prey were available in higher relative abundances.

Potential keystone OTUs in co-occurrence networks and food webs were found to be largely different (Tables 2 and 4). We identified keystones based on degree centrality, but OTUs with high closeness and between centralities were largely very similar (Appendix Tables A3.1 and A3.2). The three-spined stickleback and long-spined bullhead were the only two OTUs that exhibited high degree centrality across all four co-occurrence networks (Tables 2 & 4). The functional role of these species in ecosystems is not well understood, especially non-trophic interactions which make up the majority of interactions within our networks. However, both OTUs were most abundant in the nearshore environment and co-occurred within all networks, suggesting their co-occurrence could be driven by similar habitat requirements. The three-spined stickleback spawns in very shallow coastal environments (<3 m depth) during the spring and summer but spends most of its life cycle in open seas, whilst the long-spined bullhead is a permanent resident of the intertidal zone (Bergstrom *et al*., 2015, Barrett *et al*., 2016). These species also share lots of edges with other predominantly coastal inhabitants, such as the European herring gull (*Larus argentatus*), corkwing wrasse (*Symphodus melops*), rock gunnel (*Pholis gunnellus*), salmon/trout (*Salmo*), and many benthic invertebrate species (bivalves, gastropods, polychaetes) (Figure 2), enhancing the likelihood that co-occurrences resulted from shared habitat requirements. In this instance, these species are therefore unlikely to be keystone species with disproportionate negative effects across the ecosystem if removed (Faust et al., 2012). Other OTUs, such as bottlenose dolphins in the offshore network, and sandeels in the late season network, only have high degree centrality in the networks where they are least abundant. We sighted no bottlenose dolphins offshore whilst collecting samples, and previous research corroborates that they are regionally coastal, found in depths of less than 25 meters and typically no further than 1.5 km from the shore (Culloch & Robinson, 2008; Robinson et al., 2007). Therefore, it is likely that the small quantities of eDNA detected for bottlenose dolphins in the offshore environment resulted from the movement of DNA particles in the water column as opposed to the bottlenose dolphins actually being present (Andruszkiewicz et al., 2019). Similarly, sandeels were only detected as a keystone OTU in the late season network when they are known to occur in low abundance and are less unavailable to interact with other species in the water column since they are primarily buried in the sand (Henriksen et al., 2021). Conversely, OTUs that exert large influences in North Sea food webs, were not identified as keystone OTUs in co-occurrence networks. For example, copepods contribute up to 90% of the zooplankton biomass in the North Sea and are responsible for transferring energy from primary producers to commercially important fish species, as well as nutrient recycling and carbon export (Kürten et al., 2013; Mortelmans et al., 2021). We suspect that they fail to have significant correlations with other OTUs as a result of being present in such high abundance across all samples (Zurell et al., 2018). These examples highlight some of the potential flaws associated with both eDNA metabarcoding and correlative relationships, underscoring the need for more robust validation in our interpretation of co-occurrence networks.

In conclusion, co-occurrence analyses coupled with eDNA metabarcoding data hold great potential to improve our understanding of the status and functioning of ecosystems, including identifying species interactions and potential bio-indicator species. Here, co-occurrence networks and food webs revealed similar trends in ecosystem complexity, despite interactions forming these networks being largely different making the outcomes challenging to interpret. Trophic interactions formed a small proportion of the established co-occurrences leading to key food web components such as forage fish and copepods not being highly connected in co-occurrence networks. Therefore, we strongly recommend that co-occurrence networks should be employed alongside validation methods, such as ground truthing within well-studied ecosystems as demonstrated in this study. In this scenario, food webs and co-occurrence networks are complementary, and co-occurrence networks highlighted key taxa to focus diet analyses on to overcome current limitations, especially in planktonic communities. Further, limitations of both eDNA metabarcoding and correlation methods need to be explicitly accounted for in these analyses, as both may impact upon which interactions are detected as demonstrated in the present examination. We encourage future research exploring eDNA metabarcoding studies across different spatiotemporal scales to optimise the detection of species interactions within co-occurrence networks. These ecosystem-level analyses could then provide an early warning system for detecting ecosystem changes in response to climate warming or human pressures and highlight species or mechanisms underpinning the changes which can contribute to management or policy decisions.

## Acknowledgements

EB was funded by the Leeds Doctoral Scholarship from the University of Leeds. We would like to thank all the Cetacean Research and Rescue volunteers that participated in sample collection. We would like to thank Morag Taylor for her support with sample library preparation and sequencing. We would like to express our appreciation to Dr Elena Valsecchi for additional PhD supervision and fruitful eDNA discussions throughout the development of this manuscript.

## References

Adams, J., & Martin, J. (1986). The hydrography and plankton of the Moray Firth. *Proceedings of the Royal Society of Edinburgh*, Section B: Biological Sciences, 91, 37–56.

Amaral-Zettler, L. A., McCliment, E. A., Ducklow, H. W., & Huse, S. M. (2009). A method for studying protistan diversity using massively parallel sequencing of V9 hypervariable regions of small-subunit ribosomal RNA genes. PloS one, 4(7), e6372.

Andruszkiewicz, E. A., Koseff, J. R., Fringer, O. B., Ouellette, N. T., Lowe, A. B., Edwards, C. A., & Boehm, A. B. (2019). Modeling environmental DNA transport in the coastal ocean using Lagrangian particle tracking. Frontiers in Marine Science, 6, 477.

Araújo, M. B., & Rozenfeld, A. (2014). The geographic scaling of biotic interactions. Ecography, 37(5), 406–415.

Assenov, Y., Ramírez, F., Schelhorn, S.-E., Lengauer, T., & Albrecht, M. (2008). Computing topological parameters of biological networks. Bioinformatics, 24(2), 282–284.

Barroso-Bergada, D., Pauvert, C., Vallance, J., Deliere, L., Bohan, D. A., Buee, M., & Vacher, C. (2021). Microbial networks inferred from environmental DNA data for biomonitoring ecosystem change: Strengths and pitfalls. Molecular ecology resources, 21(3), 762–780.

Benjamini, Y., & Hochberg, Y. (1995). Controlling the false discovery rate: a practical and powerful approach to multiple testing. Journal of the Royal statistical society: series B (Methodological), 57(1), 289–300.

Berry, D., & Widder, S. (2014). Deciphering microbial interactions and detecting keystone species with co-occurrence networks. Frontiers in microbiology, 5, 219.

Blanchet, F. G., Cazelles, K., & Gravel, D. (2020). Co-occurrence is not evidence of ecological interactions. Ecology Letters, 23(7), 1050–1063.

Boettiger, C., Lang, D. T., & Wainwright, P. (2012). rfishbase: exploring, manipulating and visualizing FishBase data from R. Journal of fish biology, 81(6), 2030–2039.

Bossier, S., Nielsen, J. R., & Neuenfeldt, S. (2020). Exploring trophic interactions and cascades in the Baltic Sea using a complex end-to-end ecosystem model with extensive food web integration. Ecological modelling, 436, 109281.

Boyse, E., Robinson, K. P., Beger, M., Carr, I. M., Taylor, M., Valsecchi, E., & Goodman, S. J. (2023). Environmental DNA reveals fine scale spatial and temporal variation of prey species for marine mammals in a Scottish marine protected area. bioRxiv, 2023.2012. 2021.572838.

Boyse, E., Robinson, K. P., Beger, M., Carr, I. M., Taylor, M., Valsecchi, E., & Goodman, S. J. (2024). Environmental DNA reveals fine scale spatial and temporal variation of prey species for marine mammals in a Scottish marine protected area. Environmental DNA. Under Review.

Brown, M. B. (1975). 400: A method for combining non-independent, one-sided tests of significance. Biometrics, 987–992.

Cazelles, K., Araújo, M. B., Mouquet, N., & Gravel, D. (2016). A theory for species co-occurrence in interaction networks. Theoretical Ecology, 9, 39–48.

Csardi, G., & Nepusz, T. (2006). The igraph software package for complex network research. InterJournal, complex systems, 1695(5), 1–9.

Culloch, R. M., & Robinson, K. P. (2008). Bottlenose dolphins using coastal regions adjacent to a Special Area of Conservation in north-east Scotland. Journal of the Marine Biological Association of the United Kingdom, 88(6), 1237–1243.

D’Alessandro, S., & Mariani, S. (2021). Sifting environmental DNA metabarcoding data sets for rapid reconstruction of marine food webs. Fish and Fisheries, 22(4), 822–833.

Delmas, E., Besson, M., Brice, M. H., Burkle, L. A., Dalla Riva, G. V., Fortin, M. J., Gravel, D., Guimarães Jr, P. R., Hembry, D. H., & Newman, E. A. (2019). Analysing ecological networks of species interactions. Biological Reviews, 94(1), 16–36.

Di Muri, C., Lawson Handley, L., Bean, C. W., Benucci, M., Harper, L. R., James, B., Li, J., Winfield, I. J., & Hänfling, B. (2022). Spatio-temporal monitoring of lake fish spawning activity using environmental DNA metabarcoding. Environmental DNA.

Edwards, M., & Richardson, A. J. (2004). Impact of climate change on marine pelagic phenology and trophic mismatch. Nature, 430(7002), 881–884.

Engelhard, G. H., Peck, M. A., Rindorf, A., C. Smout, S., van Deurs, M., Raab, K., Andersen, K. H., Garthe, S., Lauerburg, R. A., & Scott, F. (2014). Forage fish, their fisheries, and their predators: who drives whom? ICES Journal of Marine Science, 71(1), 90–104.

Fauchald, P., Skov, H., Skern-Mauritzen, M., Johns, D., & Tveraa, T. (2011). Wasp-waist interactions in the North Sea ecosystem. PloS one, 6(7), e22729.

Faust, K., & Raes, J. (2016). CoNet app: inference of biological association networks using Cytoscape. F1000Research, 5, 1519.

Faust, K., Sathirapongsasuti, J. F., Izard, J., Segata, N., Gevers, D., Raes, J., & Huttenhower, C. (2012). Microbial co-occurrence relationships in the human microbiome. PLoS computational biology, 8(7), e1002606.

Foden, W. B., Young, B. E., Akçakaya, H. R., Garcia, R. A., Hoffmann, A. A., Stein, B. A., Thomas, C. D., Wheatley, C. J., Bickford, D., & Carr, J. A. (2019). Climate change vulnerability assessment of species. Wiley interdisciplinary reviews: climate change, 10(1), e551.

Ford, B. M., & Roberts, J. D. (2019). Evolutionary histories impart structure into marine fish heterospecific co-occurrence networks. Global Ecology and Biogeography, 28(9), 1310–1324.

Frederiksen, M., Elston, D. A., Edwards, M., Mann, A. D., & Wanless, S. (2011). Mechanisms of long-term decline in size of lesser sandeels in the North Sea explored using a growth and phenology model. Marine Ecology Progress Series, 432, 137–147.

Freilich, M. A., Wieters, E., Broitman, B. R., Marquet, P. A., & Navarrete, S. A. (2018). Species co-occurrence networks: Can they reveal trophic and non-trophic interactions in ecological communities? Ecology, 99(3), 690–699.

Garzke, J., Ismar, S. M., & Sommer, U. (2015). Climate change affects low trophic level marine consumers: warming decreases copepod size and abundance. Oecologia, 177, 849–860.

Goberna, M., & Verdú, M. (2022). Cautionary notes on the use of co-occurrence networks in soil ecology. Soil Biology and Biochemistry, 166, 108534.

Hansen, B. K., Bekkevold, D., Clausen, L. W., & Nielsen, E. E. (2018). The sceptical optimist: challenges and perspectives for the application of environmental DNA in marine fisheries. Fish and Fisheries, 19(5), 751–768.

Heath, M. R. (2005). Changes in the structure and function of the North Sea fish foodweb, 1973– 2000, and the impacts of fishing and climate. ICES Journal of Marine Science, 62(5), 847–868.

Henriksen, O., Rindorf, A., Brooks, M. E., Lindegren, M., & van Deurs, M. (2021). Temperature and body size affect recruitment and survival of sandeel across the North Sea. ICES Journal of Marine Science, 78(4), 1409–1420.

Hestetun, J. T., Bye-Ingebrigtsen, E., Nilsson, R. H., Glover, A. G., Johansen, P.-O., & Dahlgren, T. G. (2020). Significant taxon sampling gaps in DNA databases limit the operational use of marine macrofauna metabarcoding. Marine Biodiversity, 50(5), 70.

Hirano, H., & Takemoto, K. (2019). Difficulty in inferring microbial community structure based on co-occurrence network approaches. BMC bioinformatics, 20(1), 1–14.

Jiménez, M. F. S.-S., Cerqueda-García, D., Montero-Muñoz, J. L., Aguirre-Macedo, M. L., & García-Maldonado, J. Q. (2018). Assessment of the bacterial community structure in shallow and deep sediments of the Perdido Fold Belt region in the Gulf of Mexico. PeerJ, 6, e5583.

Kürten, B., Painting, S. J., Struck, U., Polunin, N. V., & Middelburg, J. J. (2013). Tracking seasonal changes in North Sea zooplankton trophic dynamics using stable isotopes. Biogeochemistry, 113, 167–187.

Lynam, C. P., Llope, M., Möllmann, C., Helaouët, P., Bayliss-Brown, G. A., & Stenseth, N. C. (2017). Interaction between top-down and bottom-up control in marine food webs. Proceedings of the National Academy of Sciences, 114(8), 1952–1957.

MacDonald, A., Speirs, D. C., Greenstreet, S. P., Boulcott, P., & Heath, M. R. (2019). Trends in sandeel growth and abundance off the east coast of Scotland. Frontiers in Marine Science, 6, 201.

Morales-Castilla, I., Matias, M. G., Gravel, D., & Araújo, M. B. (2015). Inferring biotic interactions from proxies. Trends in ecology & evolution, 30(6), 347–356.

Mortelmans, J., Aubert, A., Reubens, J., Otero, V., Deneudt, K., & Mees, J. (2021). Copepods (Crustacea: Copepoda) in the Belgian part of the North Sea: Trends, dynamics and anomalies. Journal of Marine Systems, 220, 103558.

NatureScot. (2020). Conservation and Management Advice: Southern Trench MPA.

O’Donnell, J. L., Kelly, R. P., Shelton, A. O., Samhouri, J. F., Lowell, N. C., & Williams, G. D. (2017). Spatial distribution of environmental DNA in a nearshore marine habitat. PeerJ, 5, e3044.

Paquette, A., & Hargreaves, A. L. (2021). Biotic interactions are more often important at species’ warm versus cool range edges. Ecology Letters, 24(11), 2427–2438.

Pethybridge, H., Bodin, N., Arsenault-Pernet, E.-J., Bourdeix, J.-H., Brisset, B., Bigot, J.-L., Roos, D., & Peter, M. (2014). Temporal and inter-specific variations in forage fish feeding conditions in the NW Mediterranean: lipid content and fatty acid compositional changes. Marine Ecology Progress Series, 512, 39–54.

Petitgas, P., Rijnsdorp, A. D., Dickey-Collas, M., Engelhard, G. H., Peck, M. A., Pinnegar, J. K., Drinkwater, K., Huret, M., & Nash, R. D. (2013). Impacts of climate change on the complex life cycles of fish. Fisheries Oceanography, 22(2), 121–139.

Pierce, G. J., Santos, M., Reid, R., Patterson, I., & Ross, H. (2004). Diet of minke whales Balaenoptera acutorostrata in Scottish (UK) waters with notes on strandings of this species in Scotland 1992–2002. Journal of the Marine Biological Association of the United Kingdom, 84(6), 1241–1244.

Ríos Castro, R., Costas Selas, C., Pallavicini, A., Vezzulli, L., Novoa, B., Teira Gonzalez, E. M., & Figueras, A. (2022). Co-occurrence and diversity patterns of benthonic and planktonic communities in a shallow marine ecosystem. Frontiers in Marine Science.

Risch, D., Castellote, M., Clark, C. W., Davis, G. E., Dugan, P. J., Hodge, L. E., Kumar, A., Lucke, K., Mellinger, D. K., & Nieukirk, S. L. (2014). Seasonal migrations of North Atlantic minke whales: novel insights from large-scale passive acoustic monitoring networks. Movement Ecology, 2(1), 1–17.

Robinson, K. P., Baumgartner, N., Eisfeld, S. M., Clark, N. M., Culloch, R. M., Haskins, G. N., Zapponi, L., Whaley, A. R., Weare, J. S., & Tetley, M. J. (2007). The summer distribution and occurrence of cetaceans in the coastal waters of the outer southern Moray Firth in northeast Scotland (UK). Lutra, 50(1), 19.

Robinson, K. P., MacDougall, D. A., Bamford, C. C., Brown, W. J., Dolan, C. J., Hall, R., Haskins, G. N., Russell, G., Sidiropoulos, T., & Sim, T. M. (2023). Ecological habitat partitioning and feeding specialisations of coastal minke whales (Balaenoptera acutorostrata) using a recently designated MPA in northeast Scotland. PloS one, 18(7), e0246617.

Russo, L., Bellardini, D., Zampicinini, G., Jordán, F., Congestri, R., & D’Alelio, D. (2023). From metabarcoding time series to plankton food webs: The hidden role of trophic hierarchy in providing ecological resilience. Marine Ecology, e12733.

Russo, L., Casella, V., Marabotti, A., Jordán, F., Congestri, R., & D’Alelio, D. (2022). Trophic hierarchy in a marine community revealed by network analysis on co-occurrence data. Food Webs, 32, e00246.

Santos, M., Pierce, G. J., Learmonth, J. A., Reid, R., Ross, H., Patterson, I., Reid, D., & Beare, D. (2004). Variability in the diet of harbor porpoises (Phocoena phocoena) in Scottish waters 1992– 2003. Marine Mammal Science, 20(1), 1–27.

Santos, M., Pierce, G. J., Reid, R., Patterson, I., Ross, H., & Mente, E. (2001). Stomach contents of bottlenose dolphins (Tursiops truncatus) in Scottish waters. Journal of the Marine Biological Association of the United Kingdom, 81(5), 873–878.

Sawaya, N. A., Djurhuus, A., Closek, C. J., Hepner, M., Olesin, E., Visser, L., Kelble, C., Hubbard, K., & Breitbart, M. (2019). Assessing eukaryotic biodiversity in the Florida Keys National Marine Sanctuary through environmental DNA metabarcoding. Ecology and Evolution, 9(3), 1029–1040.

Segers, F., Dickey-Collas, M., & Rijnsdorp, A. (2007). Prey selection by North Sea herring (Clupea harengus), with special reference to fish eggs. ICES Journal of Marine Science, 64(1), 60–68.

Shelton, A. O., Gold, Z. J., Jensen, A. J., D′ Agnese, E., Andruszkiewicz Allan, E., Van Cise, A., Gallego, R., Ramón-Laca, A., Garber-Yonts, M., & Parsons, K. (2023). Toward quantitative metabarcoding. Ecology, 104(2), e3906.

Sherr, E. B., & Sherr, B. F. (2007). Heterotrophic dinoflagellates: a significant component of microzooplankton biomass and major grazers of diatoms in the sea. Marine Ecology Progress Series, 352, 187–197.

Skelton, J., Cauvin, A., & Hunter, M. E. (2022). Environmental DNA metabarcoding read numbers and their variability predict species abundance, but weakly in non-dominant species. Environmental DNA.

Thurman, L. L., Barner, A. K., Garcia, T. S., & Chestnut, T. (2019). Testing the link between species interactions and species co-occurrence in a trophic network. Ecography, 42(10), 1658–1670.

Tylianakis, J. M., Laliberté, E., Nielsen, A., & Bascompte, J. (2010). Conservation of species interaction networks. Biological Conservation, 143(10), 2270–2279.

Valiente-Banuet, A., Aizen, M. A., Alcántara, J. M., Arroyo, J., Cocucci, A., Galetti, M., García, M. B., García, D., Gómez, J. M., & Jordano, P. (2015). Beyond species loss: the extinction of ecological interactions in a changing world. Functional Ecology, 29(3), 299–307.

Valsecchi, E., Arcangeli, A., Lombardi, R., Boyse, E., Carr, I. M., Galli, P., & Goodman, S. J. (2021). Ferries and environmental DNA: underway sampling from commercial vessels provides new opportunities for systematic genetic surveys of marine biodiversity. Frontiers in Marine Science, 1136.

Valsecchi, E., Bylemans, J., Goodman, S. J., Lombardi, R., Carr, I., Castellano, L., Galimberti, A., & Galli, P. (2020). Novel universal primers for metabarcoding environmental DNA surveys of marine mammals and other marine vertebrates. Environmental DNA, 2(4), 460–476.

Wanless, S., Harris, M., Redman, P., & Speakman, J. (2005). Low energy values of fish as a probable cause of a major seabird breeding failure in the North Sea. Marine Ecology Progress Series, 294, 1–8.

Young, J. W., Hunt, B. P., Cook, T. R., Llopiz, J. K., Hazen, E. L., Pethybridge, H. R., Ceccarelli, D., Lorrain, A., Olson, R. J., & Allain, V. (2015). The trophodynamics of marine top predators: current knowledge, recent advances and challenges. Deep Sea Research Part II: Topical Studies in Oceanography, 113, 170–187.

Zamkovaya, T., Foster, J. S., de Crécy-Lagard, V., & Conesa, A. (2021). A network approach to elucidate and prioritize microbial dark matter in microbial communities. The ISME journal, 15(1), 228–244.

Zamora-Terol, S., Novotny, A., & Winder, M. (2020). Reconstructing marine plankton food web interactions using DNA metabarcoding. Molecular ecology, 29(17), 3380–3395.

Zurell, D., Pollock, L. J., & Thuiller, W. (2018). Do joint species distribution models reliably detect interspecific interactions from co-occurrence data in homogenous environments? Ecography, 41(11), 1812–1819.

