## Supplementary Information for "Inferring species interactions from co-occurrence networks with environmental DNA metabarcoding data in a coastal marine food-web"

| | |
|---|---|
| **Supplementary Figure 1** | Positive and negative co-occurrences in the early season. |
| **Supplementary Figure 2** | Positive and negative co-occurrences in the late season. |
| **Supplementary Figure 3** | Positive and negative co-occurrences nearshore. |
| **Supplementary Figure 4** | Positive and negative co-occurrences offshore. |
| **Supplementary Table 1** | Ten OTUs with highest closeness centrality. |
| **Supplementary Table 2** | Ten OTUs with highest betweenness centrality. |

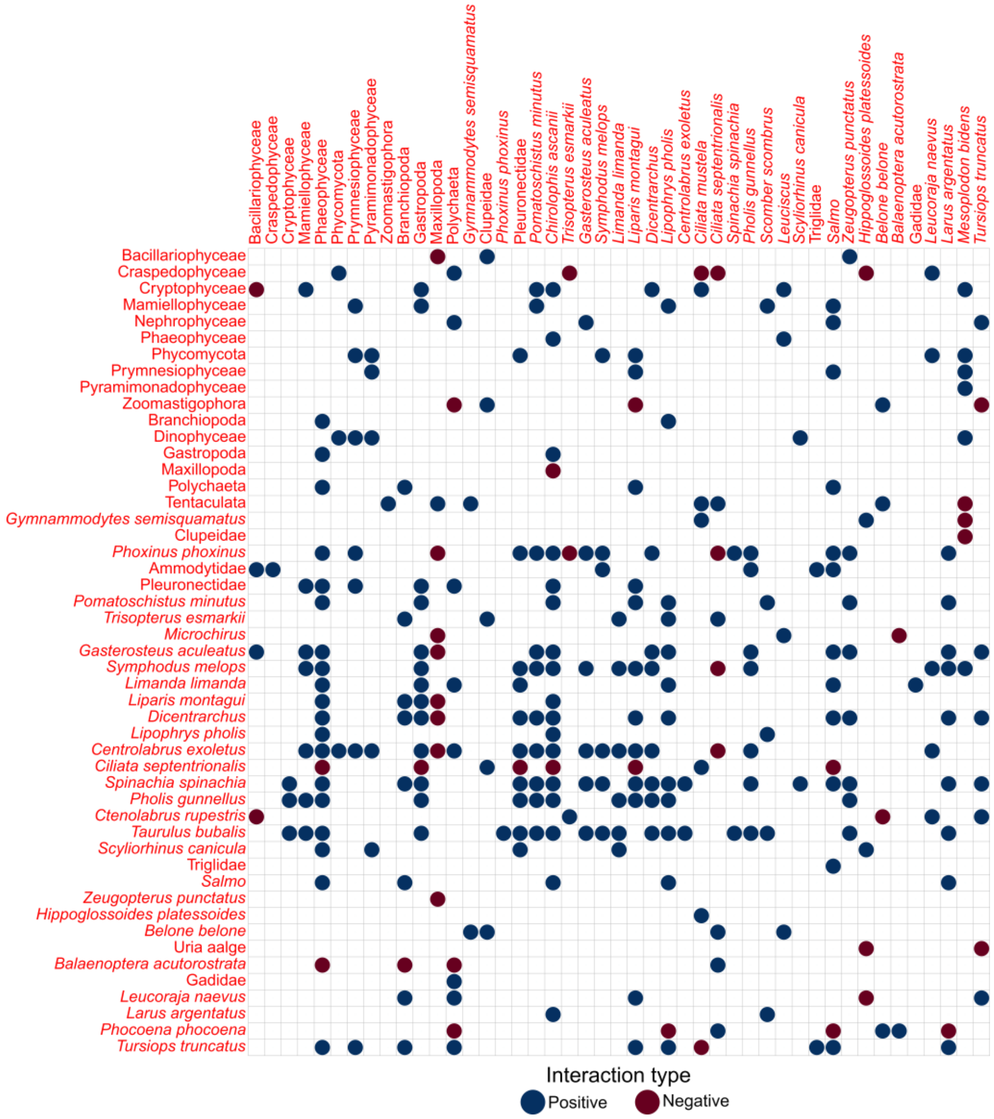


Supplementary Figure 1. Correlation matrix showing positive and negative interactions detected in the early season (June-July) co-occurrence network.

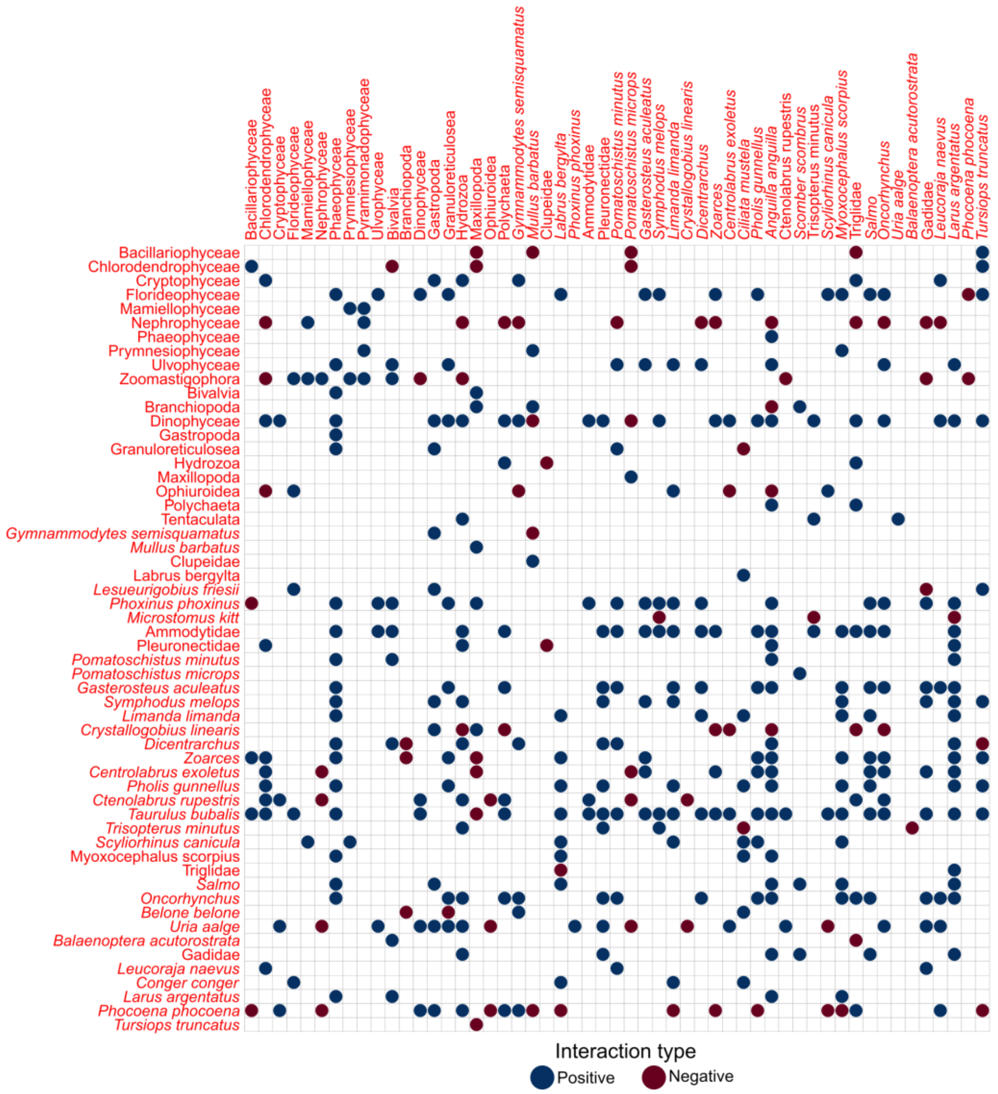


Supplementary Figure 2. Correlation matrix showing positive and negative interactions detected in the late season (August-October) co-occurrence network.

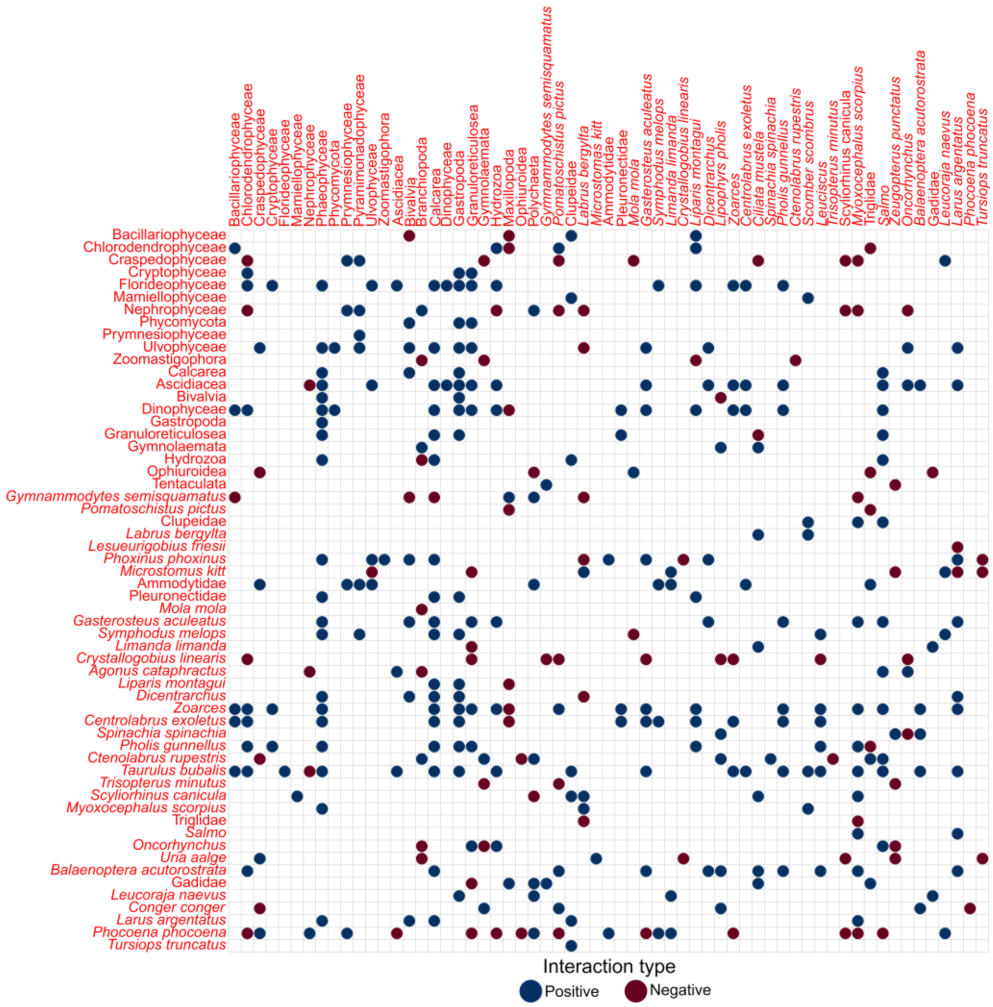


Supplementary Figure 3. Correlation matrix showing positive and negative interactions detected in the nearshore co-occurrence network.

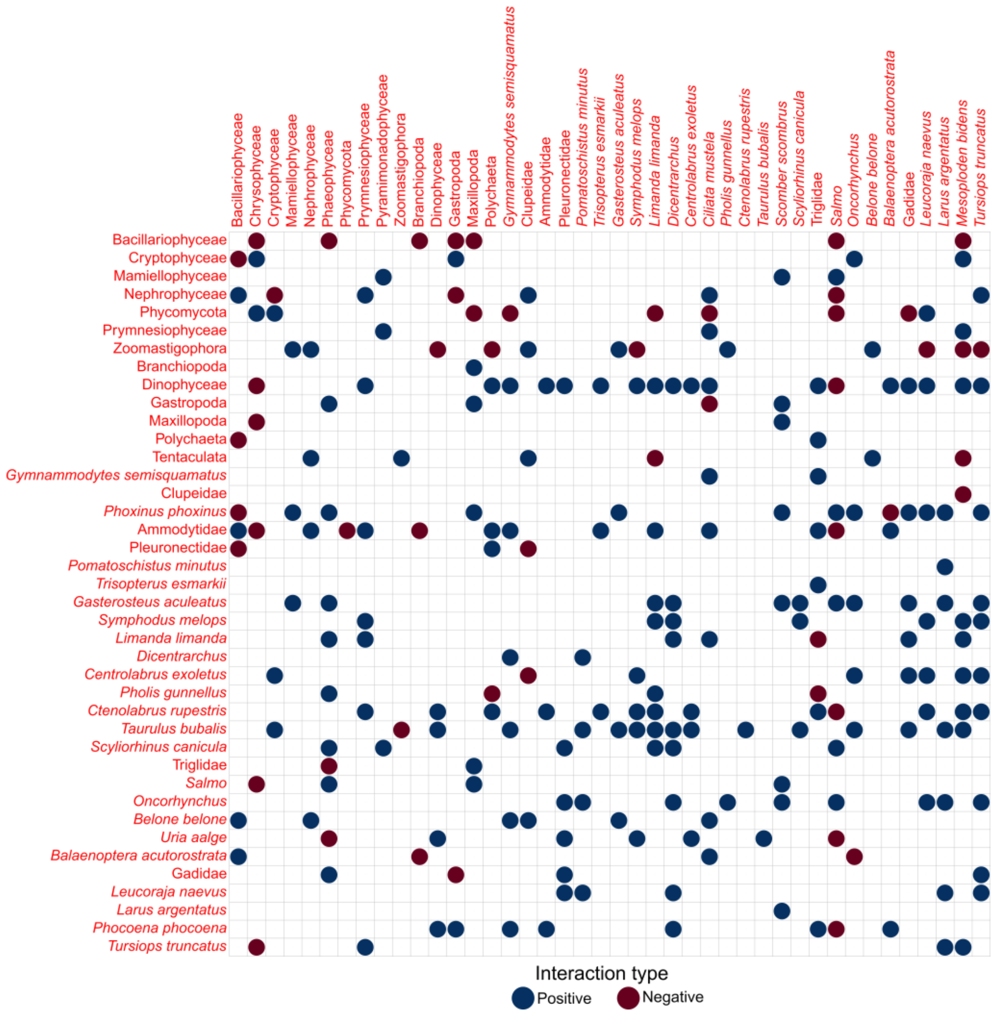


Supplementary Figure 4. Correlation matrix showing positive and negative interactions detected in the offshore co-occurrence network.

Supplementary Table 1. Ten OTUs with the highest closeness centrality in co-occurrence networks.

| Co-occurrence networks | | Co-occurrence networks | |
|---|---|---|---|
| Nearshore | Offshore | Jun-Jul | Aug-Oct |
| Zoarces | Dinophyceae | Phaeophyceae | *Oncorhynchus* |
| Calcarea | *Limanda limanda* | *Symphodus melops* | Dinophyceae |
| Ascidacea | *Salmo* | *Centrolabrus exoletus* | *Taurulus bubalis* |
| Granuloreticulosea | *Taurulus bubalis* | *Liparis montagui* | Ammodytidae |
| *Phocoena phocoena* | Ammodytidae | *Tursiops truncatus* | Zoarces |
| *Gasterosteus aculeatus* | *Tursiops truncatus* | *Salmo* | *Anguilla anguilla* |
| Ulvophyceae | *Gasterosteus aculeatus* | Dicentrachus | *Larus argentatus* |
| *Myoxocephalus scorpius* | *Oncorhynchus* | Spinachia spinachia | *Pholis gunnellus* |
| Chlorodendrophyceae | Bacillariophyceae | *Gasterosteus aculeatus* | *Phocoena phocoena* |
| Craspedophyceae | *Ctenolabrus rupestris* | *Chirolophis ascanii* | *Gasterosteus aculeatus* |

Supplementary Table 2. Betweenness centrality in co-occurrence networks.

| Co-occurrence networks | | Co-occurrence networks | |
|---|---|---|---|
| Nearshore | Offshore | Jun-Jul | Aug-Oct |
| Craspedophyceae | Dinophyceae | *Tursiops truncatus* | Dinophyceae |
| *Phocoena phocoena* | *Salmo* | *Ciliata septentrionalis* | Nephrophyceae |
| Granuloreticulosea | Ammodytidae | Phaeophyceae | Hydrozoa |
| Zoarces | *Limanda limanda* | *Salmo* | *Phocoena phocoena* |
| Larus argentatus | Bacillariophyceae | Polychaeta | *Uria aalge* |
| *Balaenoptera acutorostrata* | *Taurulus bubalis* | *Centrolabrus exoletus* | Florideophyceae |
| Ulvophyceae | *Oncorhynchus* | *Symphodus melops* | *Oncorhynchus* |
| *Myoxocephalus scorpius* | Zoomastigophora | *Liparis montagui* | *Taurulus bubalis* |
| Nephrophyceae | *Gasterosteus aculeatus* | *Gasterosteus aculeatus* | Gadidae |
| *Gymnammodytes semisquatamus* | *Tursiops truncatus* | Maxillopoda | *Larus argentatus* |

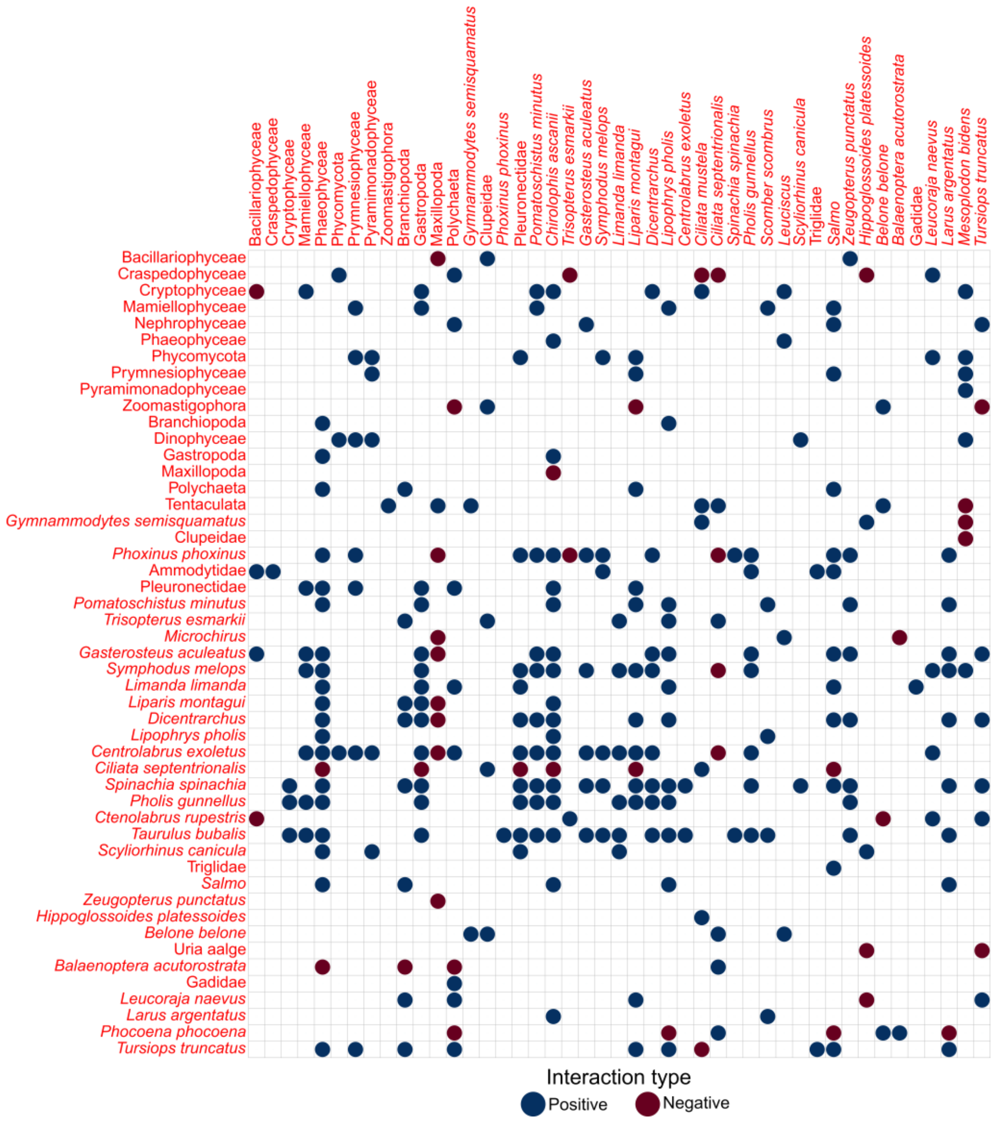


Appendix Figure A3.1. Correlation matrix showing positive and negative interactions detected in the early season (June-July) co-occurrence network.

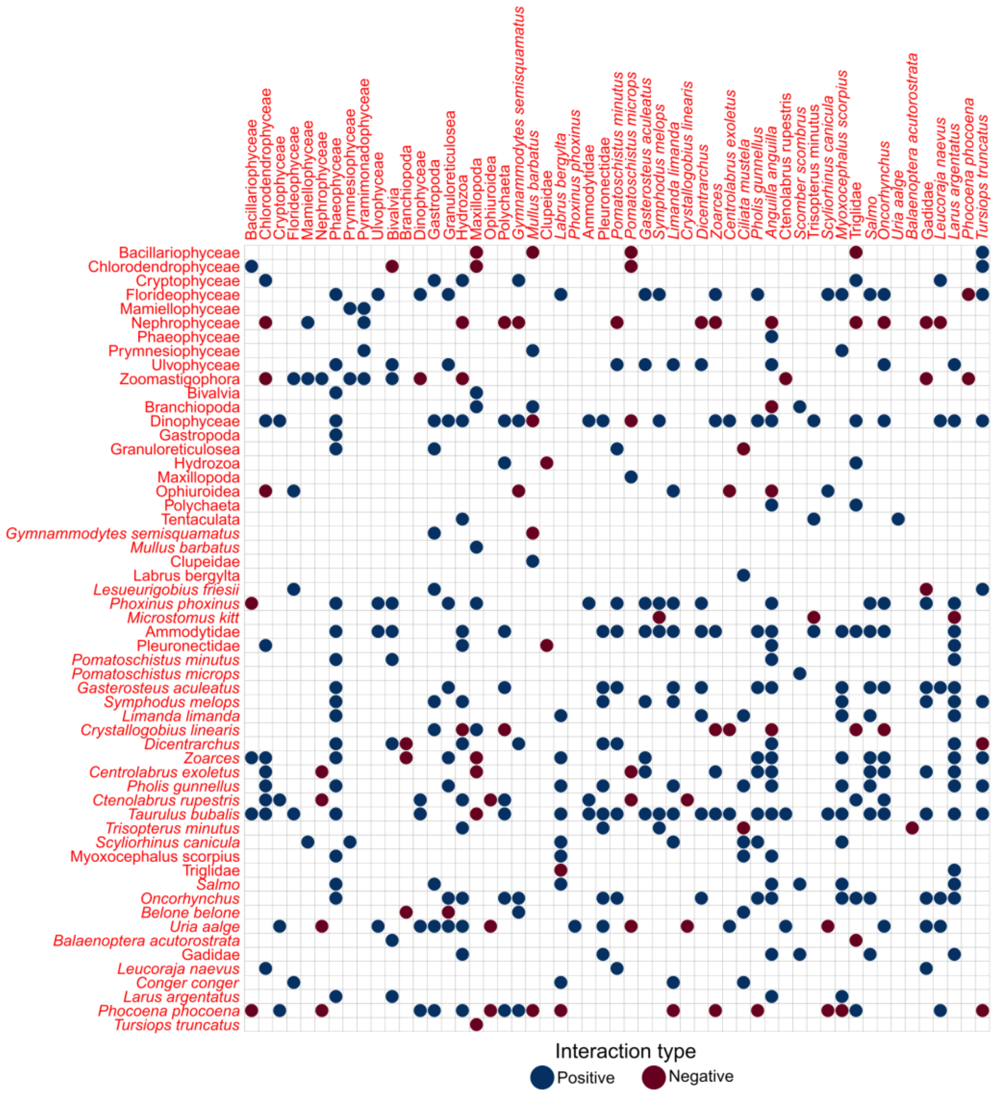


Appendix Figure A3.2. Correlation matrix showing positive and negative interactions detected in the late season (August-October) co-occurrence network.

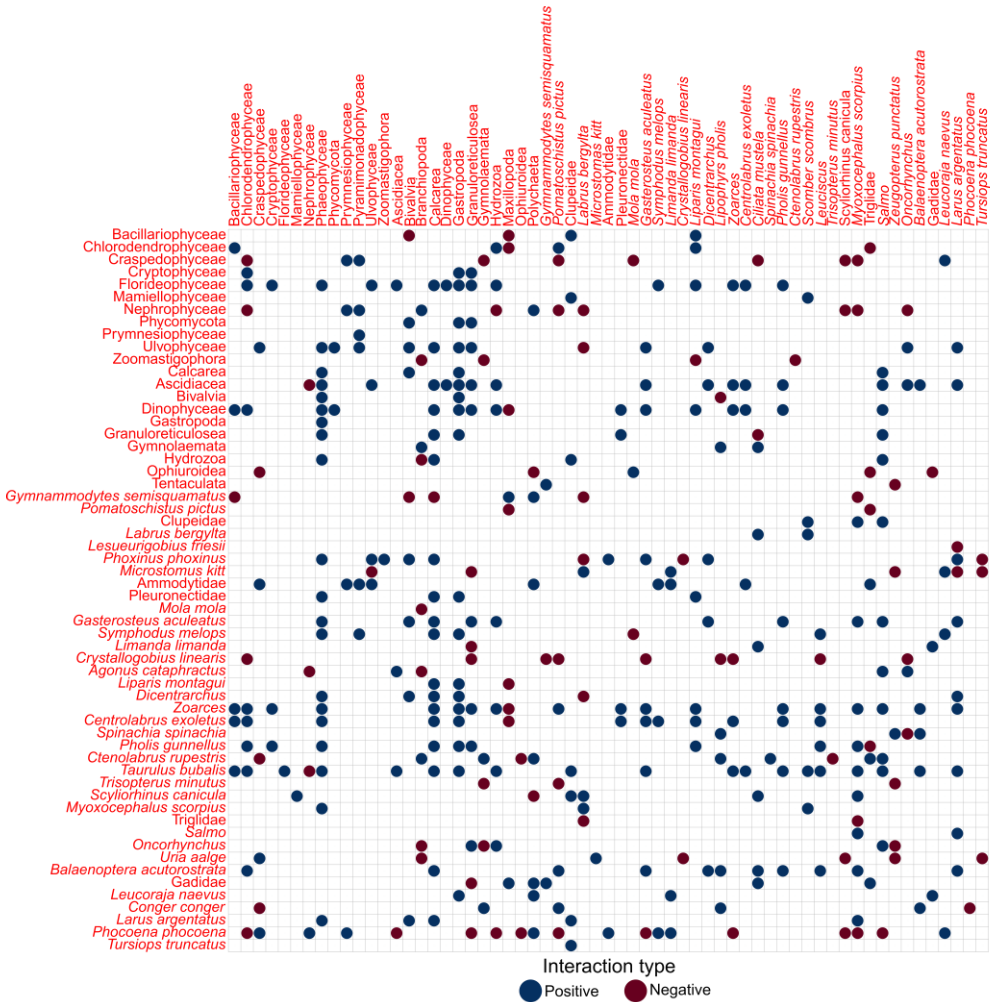


Appendix Figure A3.3. Correlation matrix showing positive and negative interactions detected in the nearshore co-occurrence network.

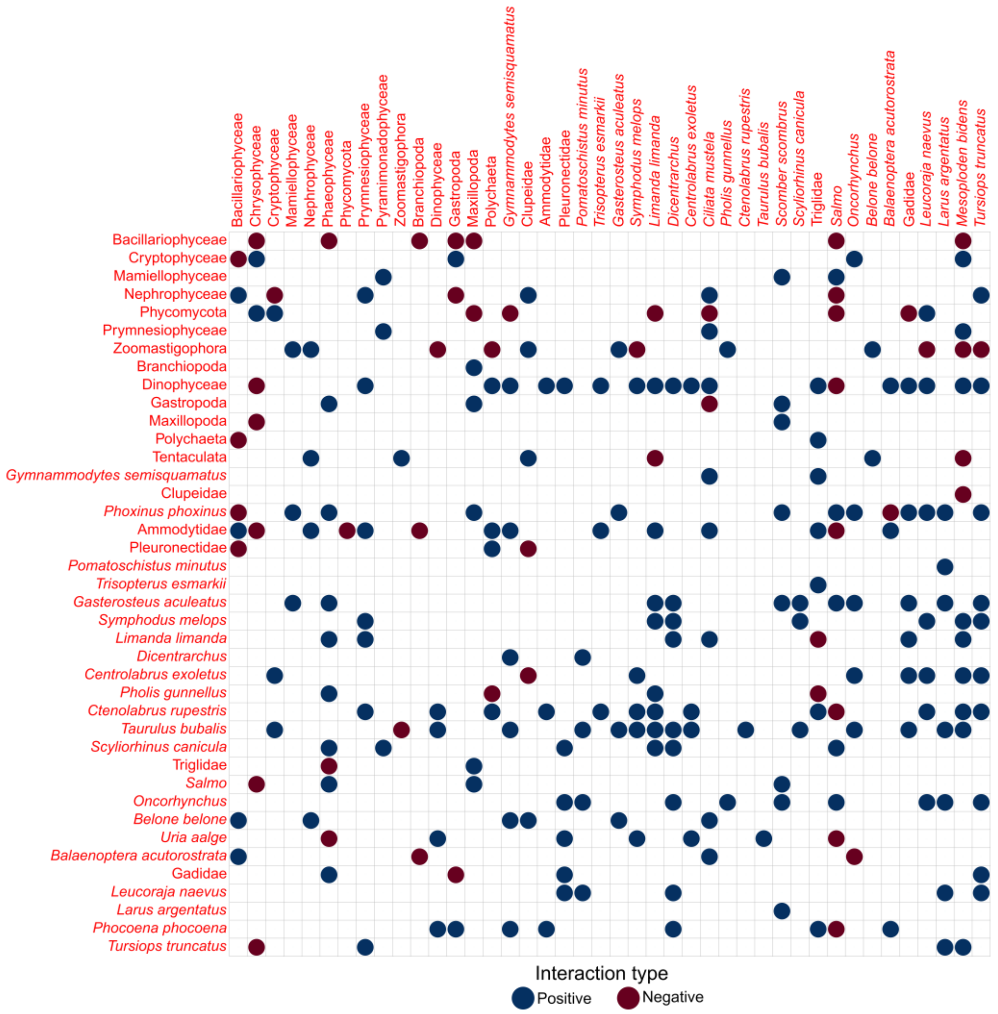


Appendix Figure A3.4. Correlation matrix showing positive and negative interactions detected in the offshore co-occurrence network.
